# MonaGO: a novel Gene Ontology enrichment analysis visualisation system

**DOI:** 10.1101/2020.09.27.316067

**Authors:** Ziyin Xin, Yujun Cai, Louis T. Dang, Hannah M.S. Burke, Jerico Revote, Hieu T. Nim, Yuan-Fang Li, Mirana Ramialison

## Abstract

MonaGO is a novel web-based visualisation system that provides an intuitive, interactive and responsive interface for performing gene ontology (GO) enrichment analysis and visualising the results. MonaGO combines dynamic clustering and interactive visualisation as well as customisation options to assist biologists in obtaining meaningful representation of overrepresented GO terms, producing simplified outputs in an unbiased manner. MonaGO supports gene lists as well as GO terms as inputs. Visualisation results can be exported as high-resolution images or restored in new sessions, allowing reproducibility of the analysis. An extensive comparison between MonaGO and 11 state-of-the-art GO enrichment visualisation tools based on 9 features revealed that MonaGO is the only platform that simultaneously allows interactive visualisation within one single output page, directly accessible through a web browser with customisable display options. In summary, MonaGO will facilitate the interpretation of GO analysis and will assist the biologists into the representation of the results.

## Background

Gene Ontology (GO)^1^ is widely used in biomedical sciences to mine large-scale datasets. GO enrichment is one of the most popular post-omics analyses for datasets generated by genomics, transcriptomics, proteomics and metabolomics assays. A myriad of web-based tools or software packages are available to perform GO enrichments or classification, including the popular tools DAVID ^2^ and PANTHER ^3^.

Inappropriate use of GO enrichment analyses can result in misleading targets and waste of resources, presenting massive hurdles to biologists ^4^. For example, if several GO categories are predicted to be enriched above the statistical threshold, which of them should be displayed? Often arbitrary decisions are made such as keeping only the “top 5 most-enriched” as a figure in publications. In addition, the redundancy of GO terms due to its hierarchical nature makes visualisation of enrichment results difficult, and often “representative terms” (*e.g.* “inflammation” or “differentiation”) are arbitrarily chosen to represent broadly-related GO categories. The emerging field of visual analytics ^5^ and its increasing use in biomedicine ^6^ can bridge these challenges by harnessing human expertise to navigate the dense information typically presented in GO enrichment analyses, resulting in a meaningful representation of overrepresented GO terms.

We have developed MonaGO, a novel interactive online visualisation system for GO enrichment analysis results. MonaGO provides a coordinated interface that retains all information, yet remains intuitive, fluid, and easy to use for lay users. Therefore, MonaGO assists biologists in making informed decisions on which enriched terms should be displayed to allow a meaningful representation and interpretation of their datasets, without compromising on objectivity by arbitrarily choosing “representative terms”.

Several tools exist that provide visualisation for GO enrichment analysis results ^7–9^ but few permit on-the-fly exploration of GO terms clustering via chord diagram visualisation. The main advantages of MonaGO over these tools include (1) its intuitive and interactive interface and comprehensive interaction options, (2) the ability to manually or systematically cluster GO terms interactively, and (3) extensive input and output options.

## Implementation

MonaGO utilises a client-server architecture and it comprises two main parts: (1) a frontend client receiving inputs from users and visualising the data, and (2) a backend server responsible for processing data, querying database and producing data for visualisation. The client mainly in JavaScript and server is built in Python.

The server consists of two Python modules. The first, server.py, utilizes *Flask_1*, a stable and scalable web application development framework. Specifically, when given a list of genes, this module sends a request to DAVID to obtain GO enrichment results. It also maintains a copy of the Gene Ontology hierarchy for visualising already enriched genes. Responses from DAVID are filtered and passed to the data-processing module. In addition, visualisation from a previously saved session can be restored by uploading a previously exported file, which already contains processed data. The *sever.py* module parses the file and sends it to client for visualisation directly. Redundant server nodes were implemented using the round-robin load balancing scheme to improve multi-user responsiveness.

The second module, *dataprocess.py*, performs data processing tasks, including calculating cluster similarity, creating hierarchical clusters, and reordering clusters for visualisation. Specifically, a hierarchical clustering algorithm is employed to cluster GO terms into clusters according to one of three options: the percentage of common genes between pairs of them (Jaccard similarity), the Resnik similarity ^10^ between gene pairs, and the SimRel similarity ^11^ between gene pairs. Algorithm 1 (Figure 8) provides a more detailed description of this procedure. In this algorithm, the user inputs the type of similarity measurement and, if a semantic similarity is chosen, the way the similarity between clusters is determined (that is, whether to use the average, minimum or maximum of the similarities between each combination of terms in the clusters).

Semantic similarities are calculated using the formulas described by Schlicker and Albrecht ^12^. In order to evaluate these formulas, two databases are used; firstly, to count the number of annotations of each GO term we use UniProtKB ^13^ (updated 14/02/19). Secondly, to access the Gene Ontology hierarchy we use *go-basic.obo* (accessed 05/03/19).

The client functions as a receiver and visualisation platform. MonaGO.js serves as the main controller of functionalities. Dynamic and interactive graphics are generated using *D3.js*, a JavaScript library allowing great control over final visualisation results. Through the visual interface generated by the client, users can intuitively interact with the visualisation and download high-resolution images from MonaGO, in PDF, PNG or SVG.

## Results and Discussion

### MonaGO’s interface allows a user-friendly interactive display of GO enrichment results

MonaGO supports three different ways of data entry: (1) submitting a list of genes and using DAVID^2^, one of the most widely used programs, to perform enrichment analysis in the background, (2) submitting gene lists and associated, pre-selected enriched GO terms for visualisation directly, and (3) importing a previously exported visualisation to restore it. MonaGO’s output options (Fig. 1) include high-resolution PNG or SVG images of a chord diagram (in the main visualisation) and the ontology hierarchy of a GO term (in the details panel), as well as JSON files that store the current state of the main chord diagram which can later be imported and restored in MonaGO.

**Figure 1.**
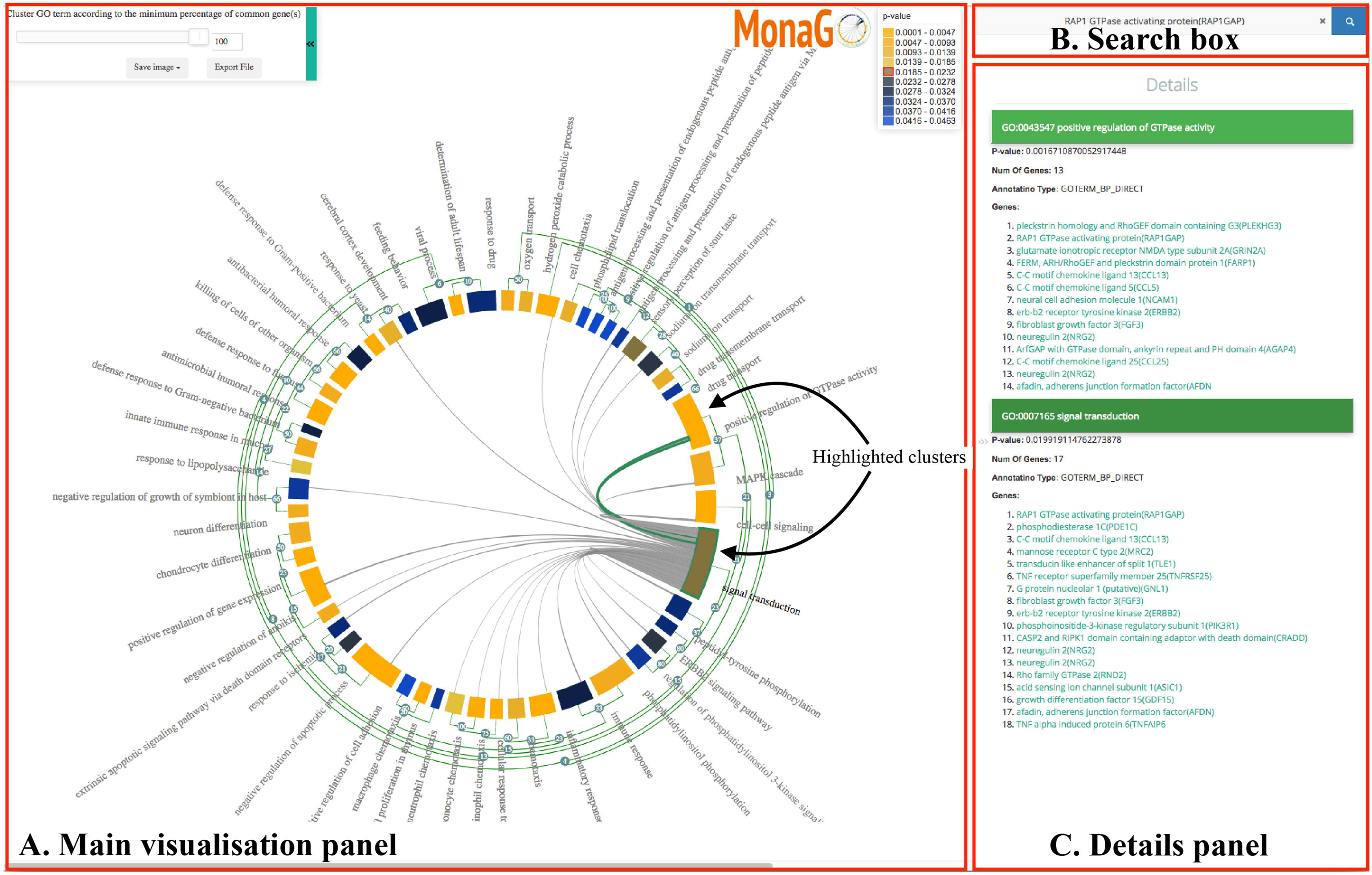
The main visualisation interface of MonaGO consisting of three components: (A, left) the main visualisation panel on the left that shows the chord diagram of GO terms that can be hierarchically and dynamically clustered, (B, top right) search box, and (C, bottom right) the details panel with dynamic GO hierarchy visualisation.

The main visualisation interface (Fig. 1) comprises three main components: (A) the main visualisation panel displays hierarchical clustering tree base on a chord diagram of enriched GO terms, (B) the “search box” panel allows browsing for specific terms or genes annotated by these terms, and (C) a “details” panel displays further information on a specific GO term upon selection.

To provide biologists with a comprehensive representation of GO enrichment results, a chord diagram (centre of Fig. 1) is employed as an intuitive and compact way to visualise clusters of GO terms and similarity between them. Enriched GO terms are colour-coded based on their p-values and their widths are proportional to the numbers of genes-of-interest contained in them.

The green edges parallel to the main chord diagram on the outside denote possible (hierarchical) clusters, and the number on an edge represents the percentage of common genes-of-interest between two nodes/clusters. The grey edges inside the chord diagram connect pairs of GO term/clusters, and the presence of such an edge denotes the existence of common genes-of-interest between them.

MonaGO helps reduce redundancy by hierarchically clustering similar GO terms in the main chord diagram. In MonaGO, enriched terms are ordered in a way that similar GO terms are placed close to each other. Users can choose between three distance metrics for the initial clustering GO terms: percentage of overlapping genes or their semantic similarity (Resnik similarity ^10^ and SimRel ^11^). When using Resnik and SimRel, users can choose between *average*, *minimum* or *maximum* options, based on the semantic similarity between each combination of individual terms in the GO clusters. *Average* takes the mean of all these similarities, and hence represents the distance between two areas of the GO tree. Alternatively, *minimum* represents the distance between the farthest two nodes of the clusters and *maximum* the closest two nodes. Hence minimum considers the Most Informative Ancestor common to all GO terms in the clusters, whereas maximum considers the Most Informative Ancestor of any two terms.

Through this chord diagram display, users can easily cluster redundant GO terms as they wish, thereby reducing the information overload. Clustering can be performed systematically or manually. A slider at top left of the main component allows systematic clustering by controlling the threshold of similarity score between terms or clusters. Alternatively, nodes/clusters in the chord diagram can be manually collapsed and expanded.

Manual clustering functionality allows users to collapse GO terms/clusters by clicking edge-nodes. For instance, as shown in Fig. 2A GO1 and GO2 share 11 common genes, which amounts to 100% of their total genes. If a user considers GO1 and GO2 to be highly redundant and wishes to group them, the green nodes between them can be clicked and thus create a cluster (Fig. 2B).

**Figure 2.**
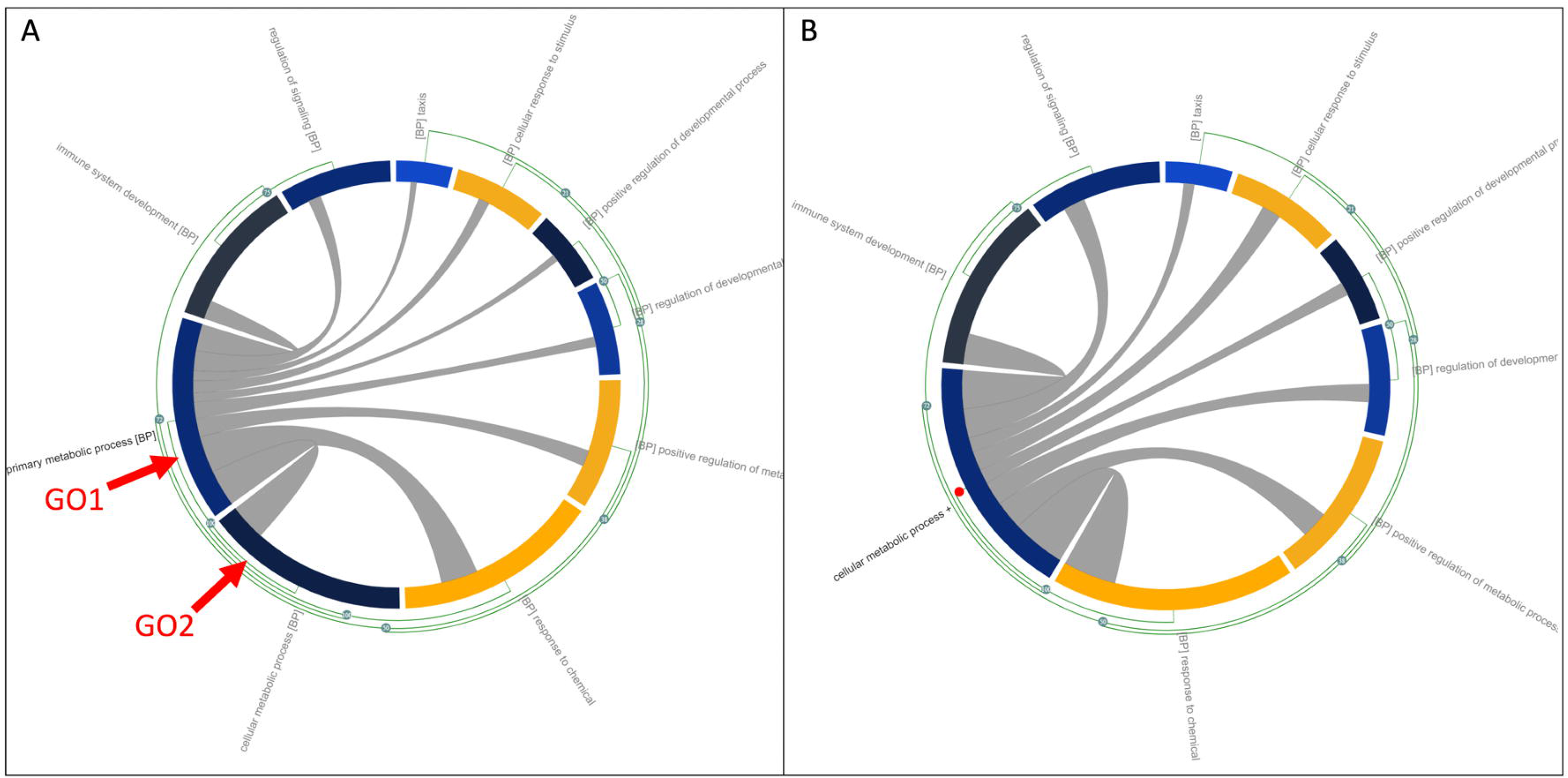
An example usage of the manual clustering feature of MonaGO: (A) the GO chord diagram before clustering where GO1 and GO2 are to be merged and, (B) the GO chord diagram after clustering.

High-resolution images of the chord diagram and the GO hierarchy of a selected GO term in the details panel can be saved in three formats, PDF, SVG and PNG, by clicking the drop-down menu button “Save image” at the top left of the main visualisation panel.

Moreover, a JSON file storing the current state of the main chord diagram and data of GO enrichment results can be downloaded and later imported into MonaGO to restore the state of the visualisation for subsequent analysis.

The details panel shows, for a node/cluster on the chord diagram or an edge inside the chord diagram, additional information about it that is complementary to the main chord diagram (Fig. 1C). For example, for an edge (highlighted with a green outline, Fig. 1A), the panel shows the number and percentage of common genes between the two nodes (GO terms) it connects to, the list of these genes and information of the semantic similarity between these terms (if chosen as similarity measure) including a diagram of the GO hierarchy (Fig. 1C). The hierarchy diagram can be expanded to full screen if needed, in order to view the tree more clearly.

Finally, the search box provides a convenient alternative way of finding genes and their associated GO terms by free-text search (Fig. 1). GO terms that correspond to a matched gene are listed in the details panel as well as popped out in the chord diagram for easy identification (*e.g.* the highlighted cluster “determination of liver left/right asymmetry in Fig. 1).

### MonaGO offers unique visualisation properties compared to existing tools

To assess MonaGO’s visualisation properties, we compared it to twelve well-known and highly-cited GO analysis systems which offer a visualisation platform (Table 1) for GO enrichment analysis. Systems such as DAVID ^2^ and PANTHER ^3^, provide a large number of analytical services while not focussing on providing an interactive visualisation interface: tables or simple graphs are used to display enrichment results. Other systems such as REVIGO ^9^, g:Profiler^15^, Gorilla ^16^, WebGestalt ^17^, and GOplot ^18^ are primarily visualisation systems dedicated to representing GO enrichment analysis results. MonaGO offers an ideal combination by providing a visualisation interface either based on existing results from GO enrichment analysis or performing GO enrichment from scratch through DAVID. Hence, where most of the systems accept GO terms (GOplot ^18^, REVIGO ^9^) or genes (WebGestalt ^17^), MonaGO offers three types of input, allowing a user to (1) submit gene lists and perform the enrichment using DAVID ^7^, (2) submitting GO terms with annotations directly, and (3) restoring previous visualisation results.

**Table 1.**
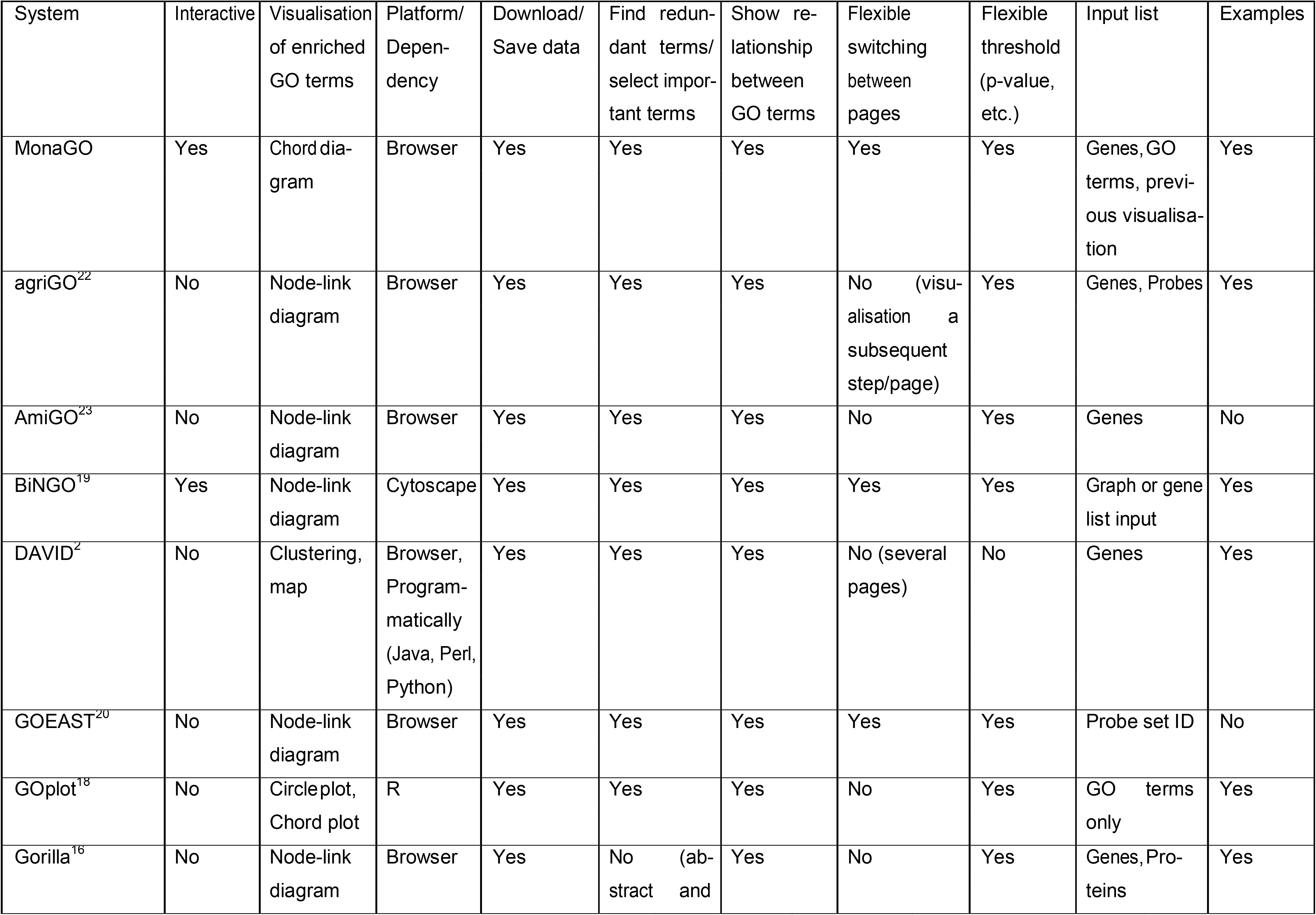
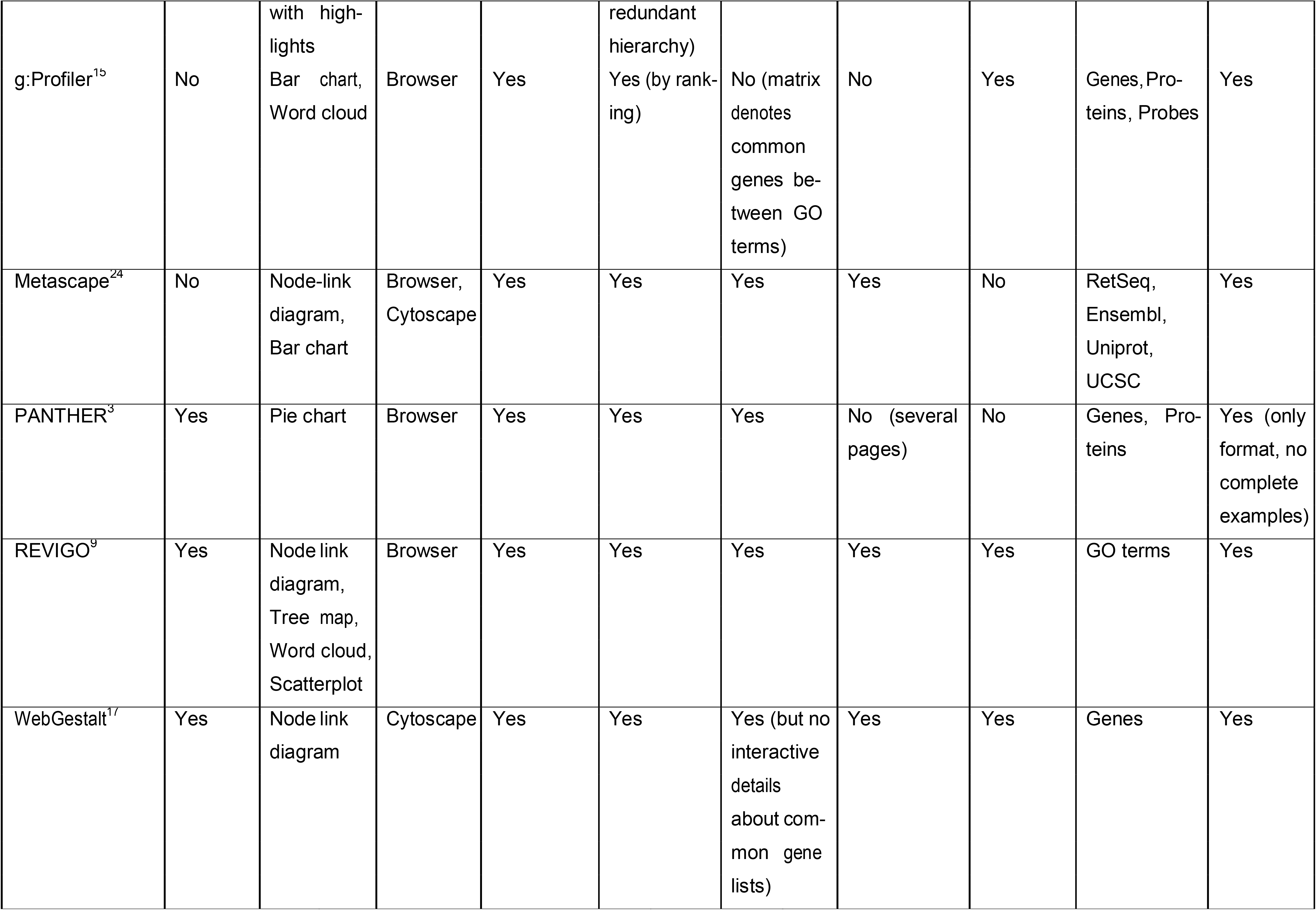
Comparisons of MonaGO with existing GO analysis systems with visualisation capabilities.

Node-link diagrams are widely used (*e.g.* BiNGO ^19^, GOEAST ^20^, Gorilla ^16^, and WebGestalt ^17^) when it comes to showing relationships between GO terms. However, the GO hierarchy or term-term relationships are not easily shown in such an approach. To address this limitation, MonaGO displays term-term similarity in a chord diagram while providing hierarchy and other information in the details panel. This novel approach allows all information to be shown while not cluttering the interface. Some tools allow modification in the visualisation output (DAVID ^2^, REVIGO ^9^, and WebGestalt ^17^) by reloading the display after re-setting the parameters of interest (such as setting threshold). Other tools (g:Profiler ^15^, Gorilla ^16^, GOplot ^18^) only provide static interfaces/images. However with MonaGO, changes in the visualisation parameters are simultaneously reflected on the output display as the user modifies them.

### MonaGO’s interactive interface facilitates the reduction of enriched GO terms to display

MonaGO is one of a few GO visualisation tools that display the relationship between terms based on the number of common genes. To illustrate the advantages of MonaGO, we re-analysed our published datasets where we measured gene expression changes for three cells types (fibroblasts, neutrophils and keratinocytes) while reprogramming the cells into a pluripotent state^21^. In brief, genes sharing similar expression levels over five stages of reprogramming were clustered using c-means fuzzy clustering. GO term enrichment for selected clusters was performed using DAVID^7^. DAVID’s default display output is a list of terms or cluster of terms (Figure 3Ai). In this test dataset, several over-represented GO terms were found enriched. Due to the length of the list, it is thus not uncommon that only the most statistically significant terms or terms relevant to the biological question are retained, creating selection bias of GO terms (Figure 3Ai). In contrast, MonaGO displays all over-represented terms (Figure 3Bi) in a single view, which can be further systematically reduced into more generic clusters (Figure 3Bii), using the overlapping number of genes as a threshold, knowledge of genes common to these clusters can be further capitalised to unravel the molecular mechanisms driving these enriched biological processes. In DAVID, this information is accessible through the cluster display mode (Figure 3Aii), where genes shared between enriched GO terms within a cluster are listed as a static heatmap. In MonaGO, at any stage during the clustering process, the genes shared between the clusters are accessible in the chord diagram (Figure 3Bii) which will assist in the interpretation of the data. For instance, in the test dataset, the most significant term “immune response” has been clustered under “immune system process”, however genes in this category are also involved in other biological processes. For example, out of 35 genes in the top category, six genes (*Ccl5*, *Tac1*,C*ccr5*, *Ccr1*, *Clec5a* and *Cd300c2*) are also involved in ‘cell-cell signaling’. MonaGO thereby allows the users to establish functional links between terms that are otherwise just presented as disjoint items in a list. Using the fibroblast dataset on REVIGO^9^, reduction of number of GO terms is effective and visualisation of these similar GO terms is clear (Figure 3Ci), based on hierarchy level and p-value. Similar clusters are retrieved through MonaGO (Figure 3Cii), however the inclusion of common genes to GO clustering provides a unique perspective on the functional relationships between GO enriched terms.

**Figure 3:**
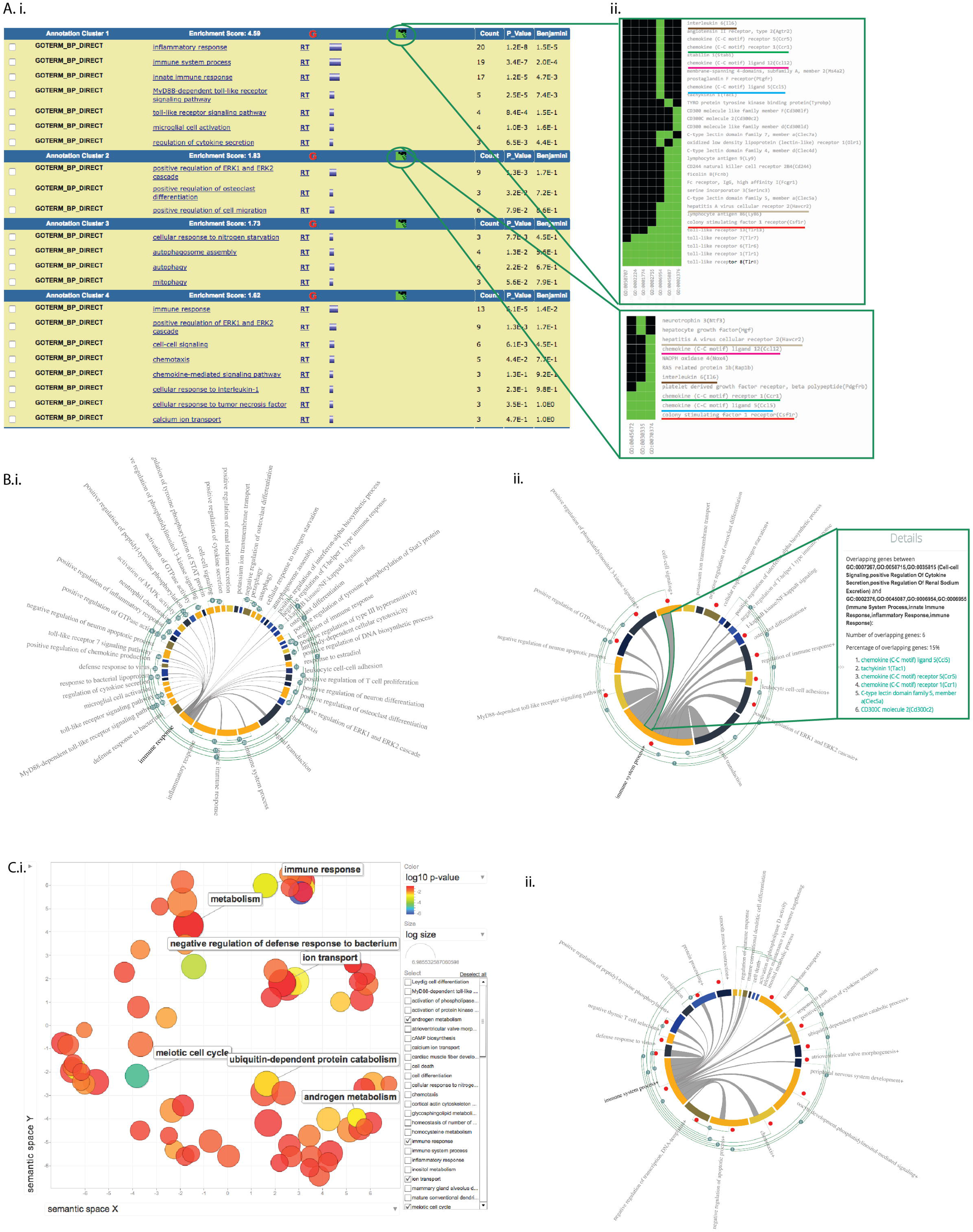
Using MonaGO to study functions of genes involved reprogramming fibroblasts to a pluripotent states. (A) List of clustered terms from GO enrichment of these genes using DAVID:(A.i) term clustering table; (A.ii) common genes display. (B) MonaGO clustering result of the same gene sets used in (A), showing (B.i) clustering of the full set of terms; and (B.ii) manual clustering from fibroblast gene cluster 4 in Nefzget et al 2017. (C) Visualisation of genes from 6 representative fibroblasts clusters by (C.i) REVIGO and (C.ii) MonaGO.

In conclusion, MonaGO‘s chord-diagram based interface allows an unbiased exploration of GO clustering results. By supporting systematic clustering of GO terms and displaying the relationships between the terms that are directly informed from the dataset, MonaGO produces meaningful representation of overrepresented GO terms in an unbiased manner.

### Clustering of GO terms by overlapping genes or semantic similarity simplifies the GO output and reveals novel functional properties

MonaGO offers two distance similarity measurement options to cluster the enriched GO terms in the chord diagram. We assessed the biological outcomes resulting from using Resnik semantic similarity versus percentage of overlapping genes, using an in-house curated list of zebrafish embryonic cardiac genes (Additional File 1). We used MonaGO to assess which biological functions compose the developmental circuitry of the heart.

Running MonaGO using “official gene symbol” as the identifier and ‘percentage of overlapping genes’ as the distance measurement allows to build a workable shortlist of biological functions that are over-represented in this gene set. As an example, running the cardiac gene set in this mode identified two different neighbouring terms ‘central nervous system projection neuron axonogenesis’ and ‘anterior/posterior axon guidance’ sharing 100% of overlapping genes (Figure 4A), hence suggesting that despite being described by different names, these two categories may represent the same function. This is further confirmed by reperforming this test using ‘Resnik similarity (average)’, where these two terms are still grouped into the same cluster (Figure 4B). Investigation of the GO hierarchy shared between the terms, which is also a feature of MonaGO, explains that their functional similarity pertains to ‘axonogenesis’ (Figure 5). Hence biologists can be confident that grouping the terms into a single node is a valid operation which also helps reduce information repetition.

**Figure 4:**
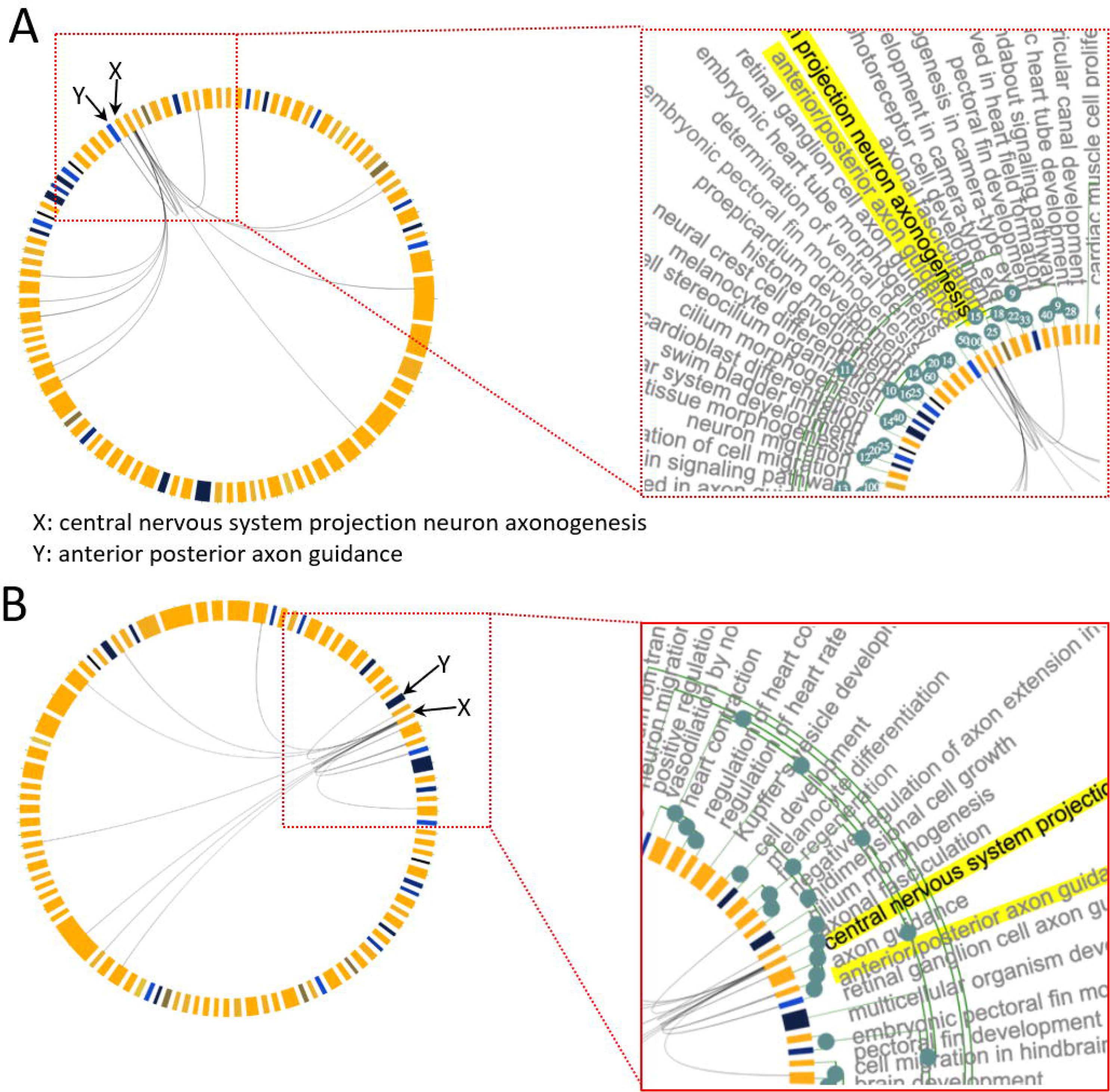
(A) Section of MonaGO’s visualisation with set of cardiac genes from zebrafish and overlapping genes as distance measurement. The GO terms ‘central nervous projection neuron axonogenesis’ and ‘anterior/posterior axon guidance’ showing 100% of overlapping genes, are highlighted in yellow. (B) Same visualisation as (A) but using Resnik similarity as distance measurement instead.

**Figure 5:**
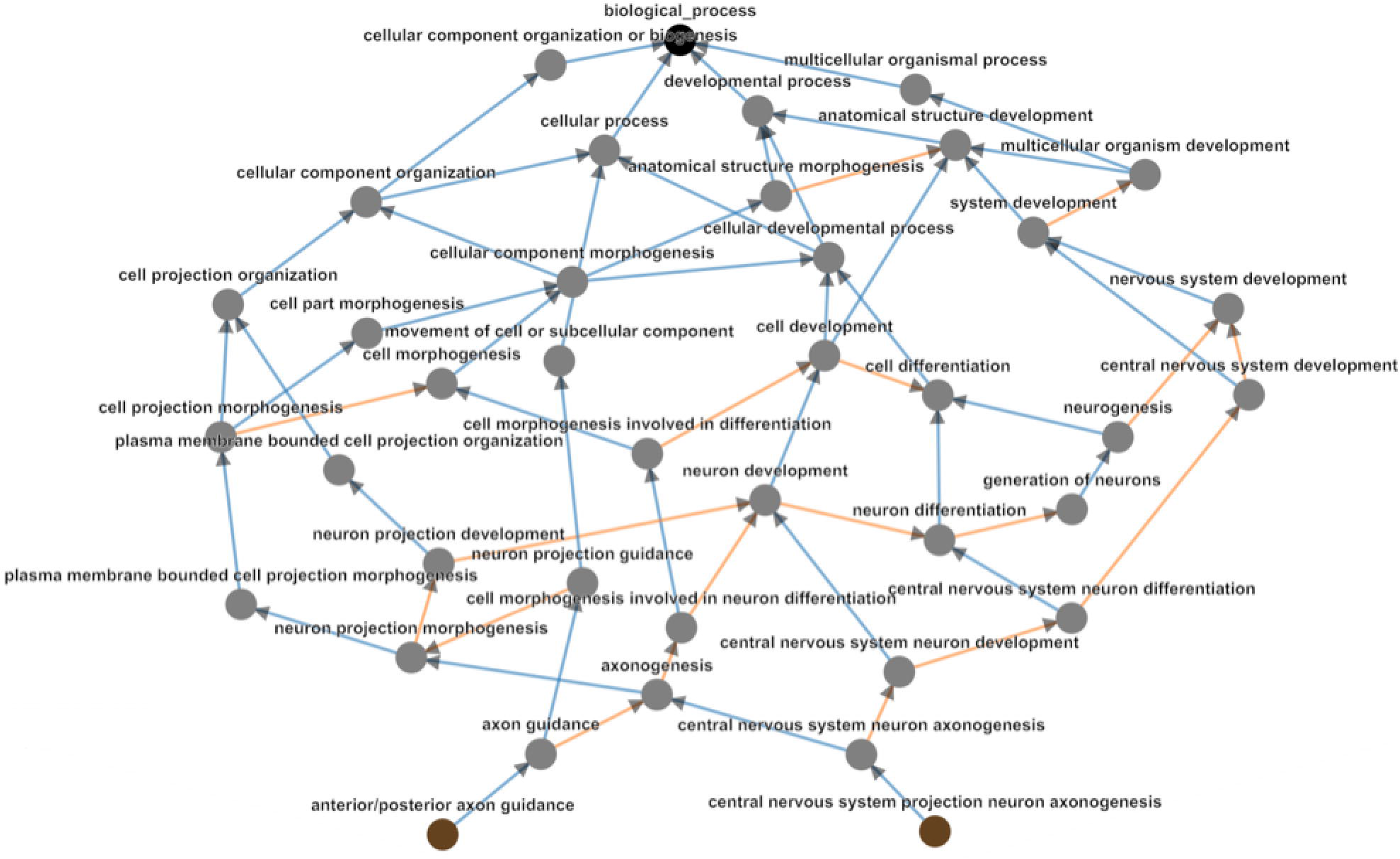
GO hierarchy between the terms ‘central nervous system project neuron axonegenesis and ‘anterior/posterior axon guidance.’ These have Resnik similarity of 3.638, with their Most Informative Ancestor being ‘axonogenesis.’

Running the cardiac gene set with “Resnik similarity (average)” as similarity measure revealed that some GO terms cluster together despite having no overlapping genes (Figure 6). Namely, the term ‘liver development’, ‘thyroid gland development’ and ‘determination of liver left/right asymmetry’ form a cluster even though there is no grey edge linking the neighbours. Thus, clustering by semantic similarity allowed us to identify two closely functionally related biological processes that are recruited in the formation of the heart, despite the lack of overlap in the genes sets composing these two processes.

**Figure 6:**
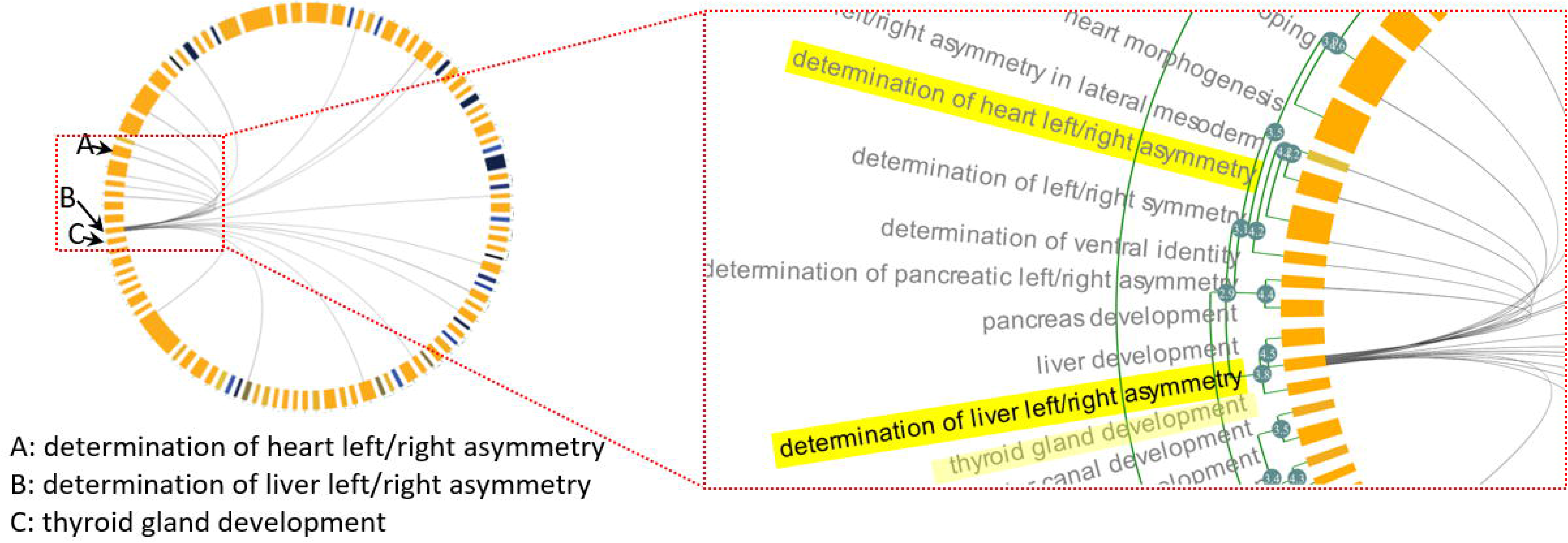
Section of MonaGO visualisation with set of cardiac genes from zebrafish and Resnik similarity as distance measurement. The GO terms ‘liver development’, ‘determination of liver left/right asymmetry’ and ‘thyroid gland development’ form a cluster of semantically similar terms with no genes overlap. This cluster shares overlapping genes with ‘determination of heart left/right asymmetry’ (highlighted in yellow).

Since this gene set is found to be active in the heart of zebrafish, we further interrogated the functional link between liver and thyroid gland development (Figure 6) and heart development. Neighbouring clusters in the chord diagram highlighted terms related to ‘left/right asymmetry’, including ‘determination of heart left/right asymmetry’. This suggests that heart, liver and thyroid gland development share common pathways during the determination of the left-right asymmetry of these organs. This common ancestor link is confirmed by the GO hierarchy (Figure 7) and supported by biological evidence as ‘liver left/right asymmetry’ and ‘determination of heart left/right asymmetry’ show 20% of overlapping genes. Most importantly, the remaining genes that belong to the liver clustered and that do not overlap with the heart cluster are of great interest for the biologists. Indeed, clustering by semantic similarity allowed them to explore a novel hypothesis that genes belonging the liver term are novel genes involved in the regulation of heart left-right asymmetry.

**Figure 7:**
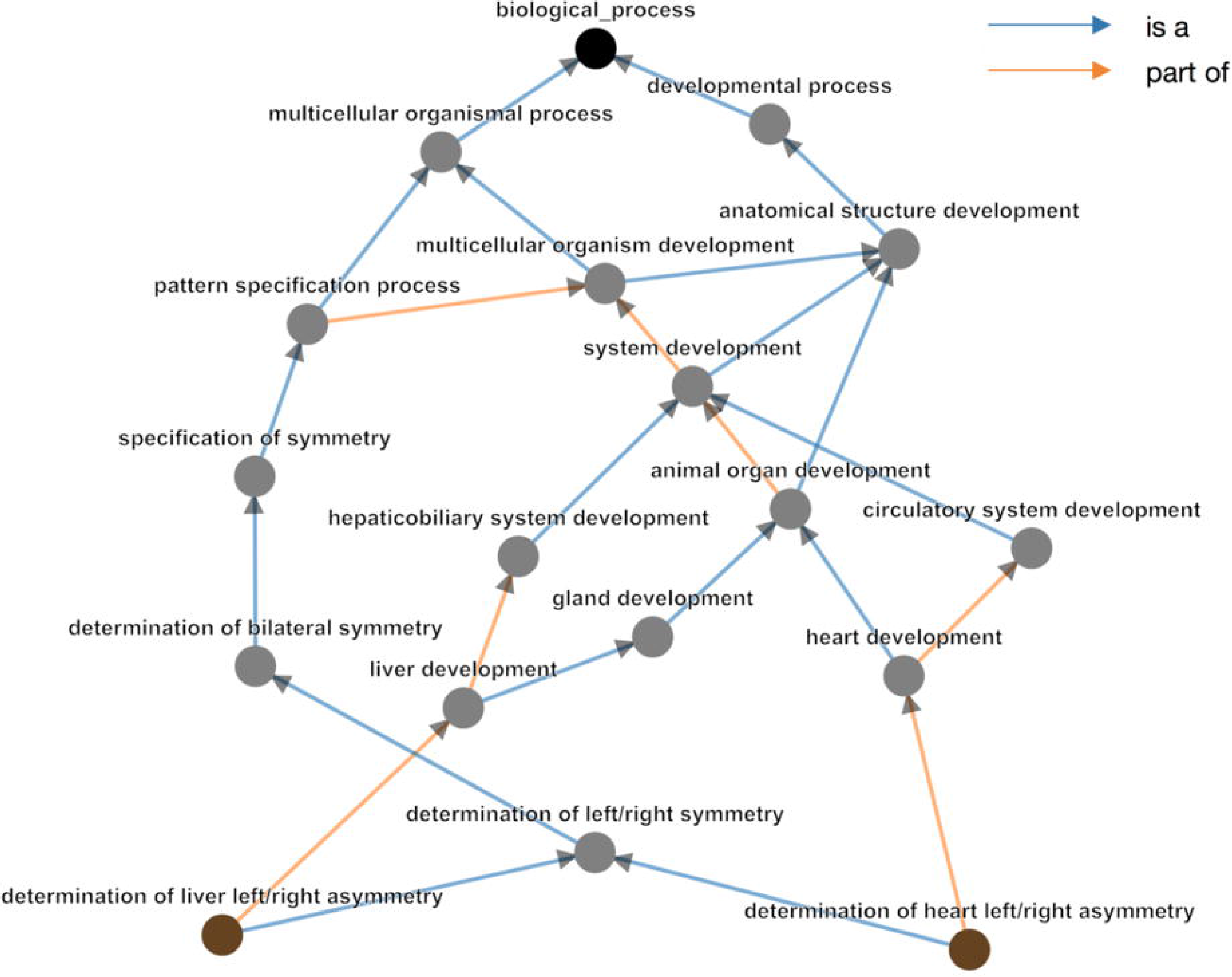
GO hierarchy between the terms ‘determination of heart left/right asymmetry’ and ‘determination of liver left/right asymmetry.’ These have Resnik similarity of 4.229, with their Most Informative Ancestor being ‘determination of left/right symmetry.’

**Figure 8:**
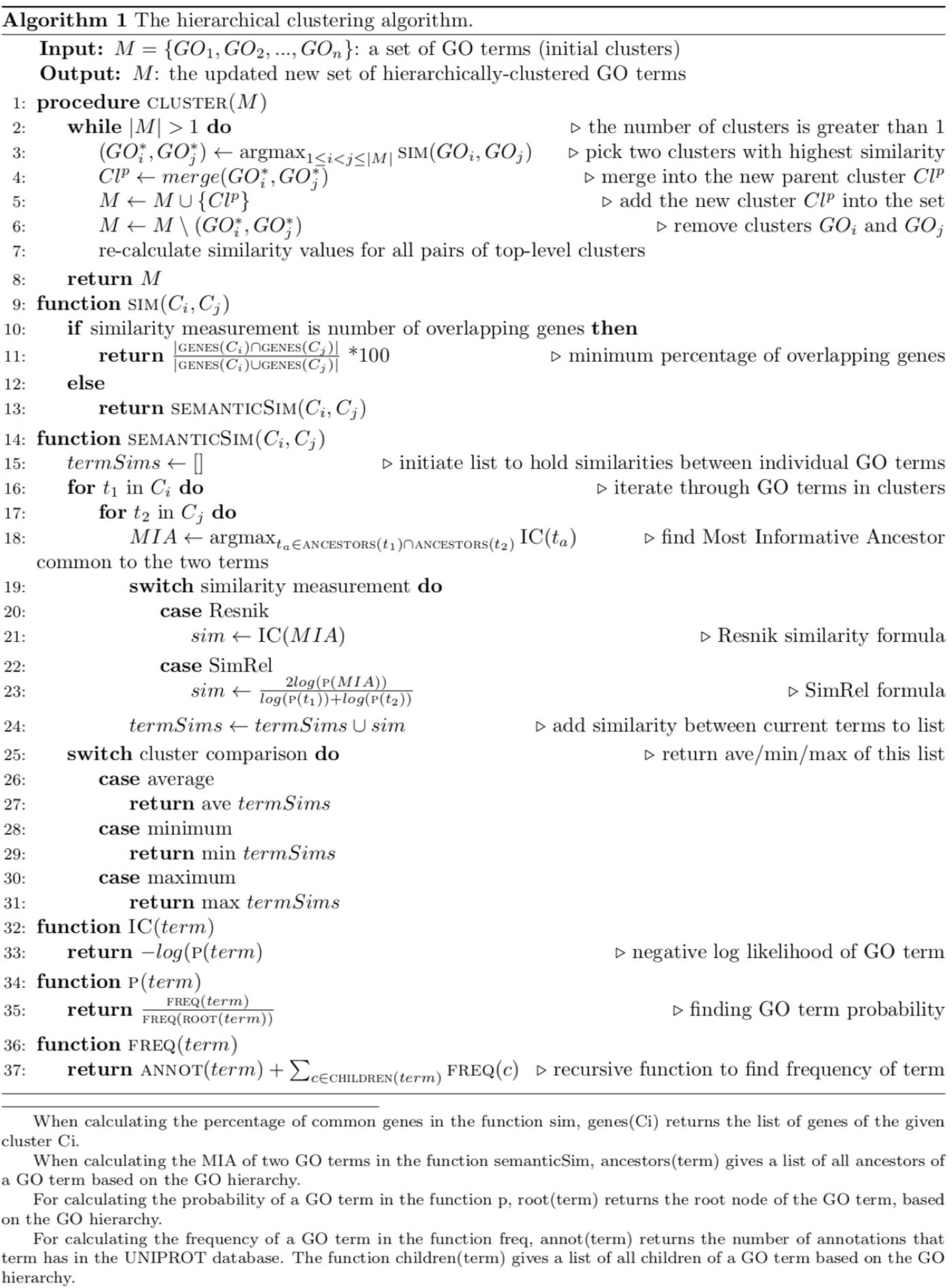
MonaGO’s hierarchical clustering algorithm (Algorithm 1) to produce the dynamic chord diagram.

## Conclusions

MonaGO is a novel web-based visualisation with unique features enabling biologists with no programming knowledge to interactive explore the GO clustering hierarchy to rapidly deduce biological interpretations. To demonstrate the benefits of MonaGO using real-world problems from developmental biologists, our platform has shown novel biological insights that may have been overlooked using traditional non-interactive exploration of the GO hierarchy. Used in combination, MonaGO’s two distance measurements provide a framework to cluster terms with optimal biological relevance and simplify the original input, even in the absence of previously known functional relationships. As a result, MonaGO aims to provide a unique tool for biologists who are interested in hands-on interaction with the gene lists and their semantic relationship to derive biological interpretation.

## DECLARATIONS

### Ethics approval and consent to participate

No ethics approval and consent required for this study.

### Consent for publication

All authors provide consent for publication.

### Authors’ contributions

M.R. and Y.-F.L. conceived the research concept and wrote the paper, Z.X. and Y.C. implemented the system and performed system comparison and analysis. L.D. provided detailed feedback to the system. HMB implemented additional distance functions for the clustering algorithm, improved the interactive visualisation, performed case studies, and revised the manuscript. HTN provided data visualisation supervision and revised the manuscript. All authors reviewed and approved the manuscript.

### Availability of data and materials

All data generated or analysed during this study are included in this published article and its supplementary information files, or directly available on the MonaGO webpage. MonaGO is freely available at http://monago.erc.monash.edu/ with all major browsers supported. The source code is available at https://github.com/liyuanfang/MonaGO.

## Acknowledgments

We thank Dr. Cristina Keightley and members of the Monash Bioinformatics Platform for their valuable feedback. We thank the Monash eResearch platform for their support with the server.

## additional files

**Additional file 1:** Curated list of embryonic cardiac genes in zebrafish

Additional_file_1.txt

